# Comparative membrane lipidomics of hepatocellular carcinoma cells reveals diacylglycerol and ceramide as key regulators of Wnt/β-catenin signaling and tumor growth

**DOI:** 10.1101/2022.08.28.505578

**Authors:** Yagmur Azbazdar, Yeliz Demirci, Guillaume Heger, Mustafa Karabicici, Gunes Ozhan

## Abstract

Hepatocellular carcinoma (HCC) is largely associated with aberrant activation of Wnt/β-catenin signaling. Nevertheless, how membrane lipid composition is altered in HCC cells with abnormal Wnt signaling remains elusive. Here, by exploiting comprehensive lipidome profiling, we unravel membrane lipid composition of six different HCC cell lines with mutations in components of Wnt/β-catenin signaling, leading to differences in their endogenous signaling activity. Among the differentially regulated lipids are diacylglycerol (DAG) and ceramide, which were downregulated at the membrane of HCC cells after Wnt3a stimulation. DAG and ceramide enhanced Wnt/β-catenin signaling in SNU475 and HepG2 cells. In contrast, depletion of DAG and ceramide suppressed Wnt/β-catenin signaling and significantly impeded the proliferation, tumor growth and *in vivo* migration capacity of SNU475 and HepG2 cells. This study, by pioneering plasma membrane lipidome profiling in HCC cells, exhibits the remarkable potential of lipids to correct dysregulated signaling pathways in cancer and stop abnormal tumor growth.

## Introduction

Hepatocellular carcinoma (HCC) is the most common form of all primary liver cancers in adults and the third common cause of cancer-related deaths worldwide. The multistage process of HCC development based on accumulation of several mutations that promote malignant transformation and tumor growth is only partially understood. Nevertheless, Wnt/β-catenin signaling pathway is known to be extensively involved in the initiation and progression of HCC (1). Owing to its key regulatory roles in development and homeostasis, Wnt signaling has been linked to cancer since the discovery that activation of *int1* (*Wnt1*) resulted in mammary hyperplasia and tumors (2–4). Wnt/β-catenin signaling is highly activated in approximately 50% of HCC cases (5, 6). Excessive Wnt signaling activation has been reported in several subgroups of HCC, i.e. the “Wnt-TGFβ subclass”, the “progenitor subclass” with mutations in AXIN1 and the relatively more heterogeneous “non-proliferation class” with mutations in the β-catenin gene CTNNB1 (7, 8). Aberrant Wnt signaling activity in HCC has been associated with enhanced tumor growth, immune escape and resistance to therapy (5,9–12).

Wnt/β-catenin (canonical Wnt) signaling is a key signaling pathway that control cell fate determination during embryonic development, maintenance of tissue homeostasis and regeneration (13–15). Wnt/β-catenin pathway is held in the inactive state in the absence of a canonical Wnt ligand, leading to phosphorylation and degradation of β-catenin by a cytoplasmic multiprotein complex, termed the β-catenin destruction complex, consisting of casein kinase 1α (Ck1α), glycogen synthase kinase 3β (Gsk3β), the tumor suppressors adenomatous polyposis coli (APC) and Axin, protein phosphatase 2A (PP2A) and the E3-ubiquitin ligase β-transducin repeats-containing protein β-TrCP (16). Signal transduction is initiated by interaction of the canonical Wnt ligand with the receptor Frizzled (Fz) and the coreceptor low-density lipoprotein-receptor-related protein 5/6 (Lrp5/6) preferentially in the ordered membrane nanodomains (17). Next, components of the β-catenin destruction complex are recruited to the Wnt-receptor complex at the plasma membrane, leading to phosphorylation of Lrp5/6 by Ck1α and Gsk3β, allowing for accumulation of β-catenin in the cytoplasm and its subsequent translocation into the nucleus, where β-catenin binds to the T-cell factor/lymphoid enhancer factor (Tcf/Lef) family of transcription factors to initiate the expression of target genes (18).

The canonical Wnt signaling pathway has been widely studied in human diseases, especially cancer, with respect to its plasma membrane components including the ligands, receptors, coreceptors and secreted or membrane-bound pathway modulators (19–22). The plasma membranes are dynamic and heterogeneous fluidic structures consisting of proteins and lipids that move via lateral, rotational and transverse diffusion, and they separate the cell from the extracellular environment (23). The membrane is further compartmentalized into numerous micro- or nanodomains that enable specialized cellular functions via regulation of receptor trafficking and downstream signaling (24). The compartmentalization can be disturbed due to mutations in genes encoding membrane proteins, abnormalities in membrane structural organization or misassembly of membrane proteins and lipids, leading to disruption of cell signaling networks and promotion of carcinogenic activities. The plasma membranes of tumor cells display notable alterations in their composition, structural organization and functional properties. Structural alterations in the membrane have also been associated with cellular multidrug resistance in cancer that may occur due to poor transport of drugs across the membrane and active efflux of therapeutic molecules from the cytoplasm to the extracellular environment via membrane transporters (25). Thus, it is rational to expect a significant difference in the membrane lipidome profiles of healthy and cancer cells (26, 27).

While membrane lipid content has been proposed to differ between cancer cells and healthy cells (28–31), membrane lipidome profiles of cancer cells, in particular in a variety of HCC cell lines with varying canonical Wnt signaling activities, have not been identified so far. This approach is especially important to test whether cancer cells can be distinguished from healthy cells with respect to their membrane lipid composition. Here, we postulate that the membrane lipidome profiles of several HCC cell lines with different endogenous Wnt signaling activities differ significantly from each other and the healthy control liver cells. To address this assumption, we took advantage of advanced shotgun lipidomics to unravel the lipids in the plasma membranes isolated from six different HCC cell lines. Initially, we validated that these HCC cells have different levels of Wnt/β-catenin signaling activity and respond differentially to Wnt pathway activation or inhibition. By globally comparing the membrane lipidome profiles of these cells, we have unraveled that their membrane lipid compositions were significantly different from each other and the healthy liver cells. Moreover, activation or inhibition of Wnt pathway in HCC cells resulted in further differentiation of their membrane lipidome profiles with respect to main lipid categories and lipid species. Among those were particular lipids that were not only differentially regulated in HCC cells but also responsive to Wnt pathway manipulation. Next, to examine whether the lipids that are differentially regulated in response to Wnt pathway manipulation can be exploited to stop tumorigenic characteristics of HCC cells, we selected the lipids diacylglycerol (DAG) and ceramide, which were both downregulated at the membrane of HCC cells after canonical Wnt pathway activation. DAG and ceramide potently activated Wnt/β-catenin signaling in SNU475 and HepG2 cells. In contrast, depletion of DAG and ceramide significantly reduced Wnt signaling and suppressed the proliferation, growth of tumor spheroids and migration of HCC cells. Overall, by unraveling the membrane lipidome profiles of HCC cells in response to manipulation of Wnt/β-catenin signaling, our study pioneers for testing the potential of cell-specific lipid fingerprint to stop tumor progression.

## Materials and Methods

### Cell culture and conditioned media production

HUH7, SNU475, Hep3B, HepG2, SNU398 and Mahlavu cells were cultured in RPMI 1640 media supplemented with 10% fetal bovine serum (FBS) at 37°C in 5% (v/v) CO2 humidified environment. THLE2 cells were grown in bronchial epithelial cell growth medium bullet kit (Lonza, Basel, Switzerland) with 10% FBS. HEK293T cells were cultured in high glucose Dulbecco’s Modified Eagle Medium (DMEM) supplemented with 10% FBS. Wnt3a was produced in murine L Wnt-3a cells (ATCC, VA, USA). L Wnt-3a cells were grown in high glucose DMEM supplemented with 10% FBS in a 100-mm tissue culture plate. When cells were 80-90% confluent, they were splitted in new 100-mm plates. Wnt3a-conditioned media (CM) were collected on the 2^nd^, 4^th^ and 6^th^ days after reaching 100% confluence and stored at 4°C until use. The secreted canonical Wnt inhibitor Dickkopf 1 (Dkk1) was produced from HEK293T cells that were seeded in a 100-mm tissue culture plate. The next day, cells were transfected with 5 µg of pCS2P+dkk1GFP plasmid in using Lipofectamine 2000 (Thermo Fisher Scientific, MA, USA). Dkk1GFP CM were collected on the 2^nd^, 4^th^ and 6^th^ days after reaching 100% confluence and stored at 4°C.

### Transfection, stimulation of cells with conditioned media and luciferase assay

Cells were seeded on 24-well cell culture plates and transfected in triplicates with 20 ng of firefly luciferase reporter pGL3 β-catenin-activated reporter (pBAR) (19099249) and 5 ng of renilla luciferase reporter pGL4.73 hRLuc/SV40 (RLuc; Promega, WI, USA) and a membrane GFP (75 ng as control for transfection) using FuGENE HD transfection reagent (1µg DNA/1µL FuGENE) (Promega, WI, USA). Next, cells were stimulated with 500 µl of Wnt3a or Dkk1GFP CM for 16 hours. Reporter activity was measured using the dual luciferase reporter assay kit (Promega, WI, USA). Statistical significance analysis was conducted using Student’s t-test. **** p < 0.0001, *** p < 0.001, ** p < 0.01, and * p < 0.05. Error bars represent standard deviation (SD).

### Isolation of giant plasma membrane vesicles (GPMVs) and lipidomics assay

Cells were seeded in 100 mm petri dishes in the appropriate media stated above, cultured to approximately 80% confluence and incubated with control, Wnt3a or Dkk1 CM for 1 hour at 37°C. Afterwards, cells were treated with 2 mM N-ethyl maleimide (NEM) in 5 mL of GMPV isolation buffer (2 mM CaCl2, 150 mM NaCl, 10 mM HEPES, pH 7.4) for 4-5 hours at 37°C (32, 33). The supernatant was transferred to a new tube and spinned at 100 g for 3 min. 4.5 mL of the supernatant was splitted into tubes and spinned at 17,000 g for 75 min at 4°C. Pellet was resuspended in 100ul of 150 mM ammonium bicarbonate. 10 ug of sample was sent for lipidomics analysis to Lipotype GmbH (Dresden, Germany).

### Lipid extraction for mass spectrometry lipidomics

Mass spectrometry-based lipid analysis was performed by Lipotype GmbH (Dresden, Germany) as described (34). Lipids were extracted using a two-step chloroform/methanol procedure (35). Samples were spiked with internal lipid standard mixture containing: cardiolipin 16:1/15:0/15:0/15:0 (CL), ceramide 18:1;2/17:0 (Cer), diacylglycerol 17:0/17:0 (DAG), hexosylceramide 18:1;2/12:0 (HexCer), lyso-phosphatidate 17:0 (LPA), lyso-phosphatidylcholine 12:0 (LPC), lyso-phosphatidylethanolamine 17:1 (LPE), lyso-phosphatidylglycerol 17:1 (LPG), lyso-phosphatidylinositol 17:1 (LPI), lyso-phosphatidylserine 17:1 (LPS), phosphatidate 17:0/17:0 (PA), phosphatidylcholine 17:0/17:0 (PC), phosphatidylethanolamine 17:0/17:0 (PE), phosphatidylglycerol 17:0/17:0 (PG), phosphatidylinositol 16:0/16:0 (PI), phosphatidylserine 17:0/17:0 (PS), cholesterol ester 20:0 (CE), sphingomyelin 18:1;2/12:0;0 (SM), triacylglycerol 17:0/17:0/17:0 (TAG). After extraction, the organic phase was transferred to an infusion plate and dried in a speed vacuum concentrator. 1st step dry extract was re-suspended in 7.5 mM ammonium acetate in chloroform/methanol/propanol (1:2:4, v:v:v) and 2nd step dry extract in 33% ethanol solution of methylamine/chloroform/methanol (0.003:5:1; v:v:v). All liquid handling steps were performed using Hamilton Robotics STARlet robotic platform with the Anti Droplet Control feature for organic solvents pipetting.

### MS data acquisition

Samples were analyzed by direct infusion on a QExactive mass spectrometer (Thermo Fisher Scientific, MA, USA) equipped with a TriVersa NanoMate ion source (Advion, NY, USA). Samples were analyzed in both positive and negative ion modes with a resolution of Rm/z=200=280000 for MS and Rm/z=200=17500 for MSMS experiments, in a single acquisition. MSMS was triggered by an inclusion list encompassing corresponding MS mass ranges scanned in 1 Da increments (36). Both MS and MSMS data were combined to monitor CE, DAG and TAG ions as ammonium adducts; PC, PC O-, as acetate adducts; and CL, PA, PE, PE O-, PG, PI and PS as deprotonated anions. MS only was used to monitor LPA, LPE, LPE O-, LPI and LPS as deprotonated anions; Cer, HexCer, SM, LPC and LPC O- as acetate adducts.

### Data analysis and post-processing

Data were analyzed with in-house developed lipid identification software based on LipidXplorer (37, 38). Data post-processing and normalization were performed using an in-house developed data management system. Only lipid identifications with a signal-to-noise ratio >5 and a signal intensity 5-fold higher than in corresponding blank samples were considered for further data analysis.

### Bioinformatics analysis

The raw lipid concentrations (in pmol) were used in data analysis. These were normalized by the sum of concentrations of all lipid species within each sample, and averaged across replicates to calculate concentration fold change between all cancer cell lines and healthy control cells, and between stimulated and nonstimulated cells of each cell line. The significance of differential regulation for each lipid was assessed by a two-tailed Student’s t-test between normalized concentrations. Missing values (i.e., non-measurable concentrations below the detection limit) were neglected in calculations, and only lipids with at least two replicates in both conditions were considered in each contrast. Statistical significance was called at p-value < 0.05. Differential regulation contrasts and lipids were clustered by Pearson correlation of log2 fold change values (missing values and non-significant effects were replaced by zero) and shown on a heatmap. Relative concentrations (in mol%/sample) were used in principal component analysis. Data was plotted using the packages ggplot2 (3.3.2), pheatmap (1.0.12), ComplexHeatmap (2.4.2), and UpSetR (1.4.0) (39–42).

### Treatment of cells with lipids and lipid synthesis inhibitors

Cells were seeded on 24-well culture plates and transfected in triplicates with 20 ng of pBAR, 5 ng of RLuc and a membrane GFP using the FuGENE HD transfection reagent as described above. 24 hours after transfection, the chemical/enzyme was added to the SNU475 cells at the concentrations indicated as follows: diacylglycerol (DAG, 50 µg/mL, Avanti polar lipid, AL, USA), ceramide (25 µg/mL, Avanti polar lipid, AL, USA) diacylglycerol kinase (DGK, 0.5 µg/mL MyBioSource, CA, USA) or myriocin (12.5 µM, Cayman Chemical Company, Michigan, USA). For HepG2 cells, the concentrations of the chemical/enzyme were as follows: DAG 200 µg/mL, ceramide 100 µg/mL, DGK 0.5 µg/mL and myriocin 12.5 µM. SNU475 and HepG2 cells were incubated at 37°C for 4 hours with DAG, ceramide and DGK and 48h with myriocin, and treated with Wnt3a CM containing the drugs for 16 hours. Reporter activity was measured using dual luciferase reporter assay kit and significance was tested using Student’s t-test as described above.

### Immunofluorescence staining

SNU475 and HepG2 cells were seeded on coverglass in a 24-well tissue culture plate. Cells were treated with DAG, ceramide and DGK for 4h or myriocin for 48h at the concentrations stated in the previous sub-section. Next, control CM, Wnt3a CM or Dkk1 CM was added into plates and incubated overnight at 37^ᵒ^C in 5% (v/v) CO_2_ humidified environment. After incubation, cover glasses were removed and cells were fixed with 4% PFA in PBS for 15 min. They were washed three times with PBS for 5 min each, PBS including 0.5% Tween-20 for 10 min, three times with PBS for 5 min each and incubated with blocking buffer (90% PBS, 1% goat serum, 0.1% Tween 20, 1% BSA, 2.25% glycine) for 30 min. Next, cells were incubated with the primary antibodies rabbit anti-phospho-β catenin (Ser675, 1:100, D2F1, Cell Signaling Technology, MA, USA) and mouse anti-β-catenin (1:100, 12F7, ab22656, Abcam, MA, USA) in PBS containing 1% BSA and 0.3% Triton-X overnight at 4^ᵒ^C. The next day, cells were washed three times with PBS for 5 min each, inbucated with the secondary antibodies goat anti-mouse IgG, superclonal recombinant secondary antibody, Alexa Fluor 488 (1:800, ThermoFisher Scientific, MA, USA) and goat anti-Rabbit IgG cross-adsorbed secondary antibody, Alexa Fluor 594 (1:800, ThermoFisher Scientific, MA, USA) for 2 hours at room temperature. Next, cells were washed five times with PBS for 5 min each, their nuclei were counterstained with 4′,6-diamidino-2-phenylindole (DAPI, 0.5 ug/ml, 4083S, Cell Signaling Technology, MA, USA) and imaged by using an LSM 880 laser scanning confocal microscope (Carl Zeiss AG, Oberkochen, Germany).

### Colony formation assay

SNU475 and HepG2 cells were seeded as 1000 cells/well in a 6-well culture plate and let grow into colonies for 8 days. After removal of the supernatant, methanol was added onto the cells and kept at - 20°C for 10 min. Next, cells were stained with crystal violet (0.5% w/v in distilled water) for 20 min at room temperature (RT) and rinsed several times with distilled water. Plates were left to dry in the incubator and imaged under a light microscope. The number of colonies were calculated by ImageJ Fiji cell counter plugin (ImageJ, U. S. National Institutes of Health, Bethesda, Maryland, USA). Significance was tested using Student’s t-test.

### Spheroid formation assay

The hanging drop method was used for this assay. Droplets of 30 µl cell culture media containing 1000 cells were pipetted on the interior of a 100 mm petri dish and the dish lid was inverted. Cells were incubated at 37°C for 4 days until spheroid formation. Next, the media was replaced with media containing the lipid synthesis inhibitors. After 24 hours or 48 hours, the cells were imaged under a stereomicroscope. Spheroid areas were calculated using ImageJ. Significance was tested using Student’s t-test.

### Zebrafish xenograft assay

SNU475 and HepG2 cells were trypsinized in cell culture plates, washed with phosphate-buffered saline (PBS) and incubated with 2mg/ml DiI in PBS at 37°C for 20 min. After washing once with FBS and twice with PBS, cells were resuspended in 10% FBS in 1X PBS at a final density of 40,000 cells/µl. Wild-type AB zebrafish larvae were dechorionated at 2 days post-fertilization (dpf) by incubating in 0.1 mg/mL pronase (Sigma-Aldrich, MO, USA) solution for 10 min at 28°C. Larvae were then anesthetized with 1 mg/mL Tricaine in E3 medium and transferred to a microinjection plate of 3% agarose in E3. Borosilicate glass capillaries with a length of 4 inches and an OD of 1.0 mm (World Precision Instruments, FL, USA) were used for microinjection. 200-300 of SNU475 cells and 700-800 of HepG2 cells were injected into the yolk sac of wild-type AB zebrafish larvae. Larvae were incubated at 34°C in fresh E3 overnight. The next day, larvae that showed infiltration of tumor cells into blood circulation were discarded. Larvae with injected cells in the yolk sac were kept and processed further for drug treatments. Zebrafish larval xenografts were fixed in 4% paraformaldehyde in PBS at 6 dpf and 7 dpf for SNU475 cells and HepG2 cells, respectively. Xenografts were imaged and the percentage of micrometastasis was determined. Significance was tested using Student’s t-test.

## Results

### Different HCC cell lines vary in their activity of Wnt/β-catenin signaling and plasma membrane lipid composition

The lipid compositions significantly vary between plasma membranes of different cell types, resulting in distinct membrane properties that in turn strongly influence the activity of membrane-bound proteins and related signalling pathways. Understanding this interdependency necessitates a good knowledge of the lipid composition of the membranes. Thus, to examine the relationship between the lipid composition of the plasma membrane and regulation of Wnt/β-catenin signaling activity, we aimed to exploit high-end lipidomics technology to unravel the membrane lipidome profiles of six different HCC cell lines. Initially, we set out to determine the endogeneous canonical Wnt signaling activities in different HCC cell lines, i.e. HUH7, SNU475, Hep3B, HepG2, SNU398 and Mahlavu cells (Figure 1A). According to the activation of the pBAR reporter of Tcf/Lef-mediated transcription, Mahlavu cells exhibited the lowest signaling activity while HepG2 and SNU398 cells had the highest activity. Signaling activities in HUH7, SNU475 and Hep3B cells were much lower than those in the HepG2 and SNU398 cells (Figure 1A). Next, to test the response of the HCC cells to manipulation of Wnt signaling, we treated the cells with the conditioned media (CM) of the canonical ligand Wnt3a or the canonical Wnt inhibitor Dkk1. Wnt3a was able to activate Wnt signaling in HUH7, SNU475 and Hep3B cells (Figure 1B). In contrast, Dkk1 stimulation could significantly suppress Wnt signaling in all five HCC cell lines (Figure 1C). Thus, to perform lipid fingerprinting we collected giant plasma membrane vesicles (GPMVs) from the cells and formed three experimental groups: i) Non-stimulated group: HUH7, SNU475, Hep3B, HepG2, SNU398, Mahlavu cells compared to the healthy control (THLE2) cells (Figure 1D). ii) Wnt3a-stimulated group: HUH7-Wnt3a, SNU475-Wnt3a and Hep3B-Wnt3a cells compared to HUH7, SNU475 and Hep3B cells (Figure 1E). iii) Dkk1-stimulated group: HepG2-Dkk1 and SNU398-Dkk1 cells compared to HepG2 and SNU398 cells (Figure 1F). Following lipid extraction and mass spectrometry-based lipid analysis performed by Lipotype GmbH, we performed principal component analysis (PCA) on the three experimental groups (Figure 1D-F). Individual samples of non-stimulated HCC cells were well separated from each other and the control samples (Figure 1D). There was a remarkable variance between healthy control positioned on one side and all non-stimulated HCC cells on the other side (Figure 1D). In particular, SNU398 and SNU475 clearly stood out from the other cells. PCA revealed that induction of HCC cells with Wnt3a or Dkk1 resulted in changes in their lipid composition and significant separation from each other (Figure 1E-F). Wnt3a stimulation resulted in minor changes on the membrane lipidome of HUH7 and SNU475 cells, whereas strong variance was detectable in Hep3B cells induced with Wnt3a (Figure 1E). Dkk1 stimulation resulted in significant changes of the membrane lipidome in both HepG2 and SNU398 cells (Figure 1F). Together, these results indicate that HCC cells with different levels of endogenous Wnt/β-catenin activity differ in their response to Wnt pathway manipulation and that this differential response of HCC cells is also reflected in their membrane lipid composition.

**Figure 1:**
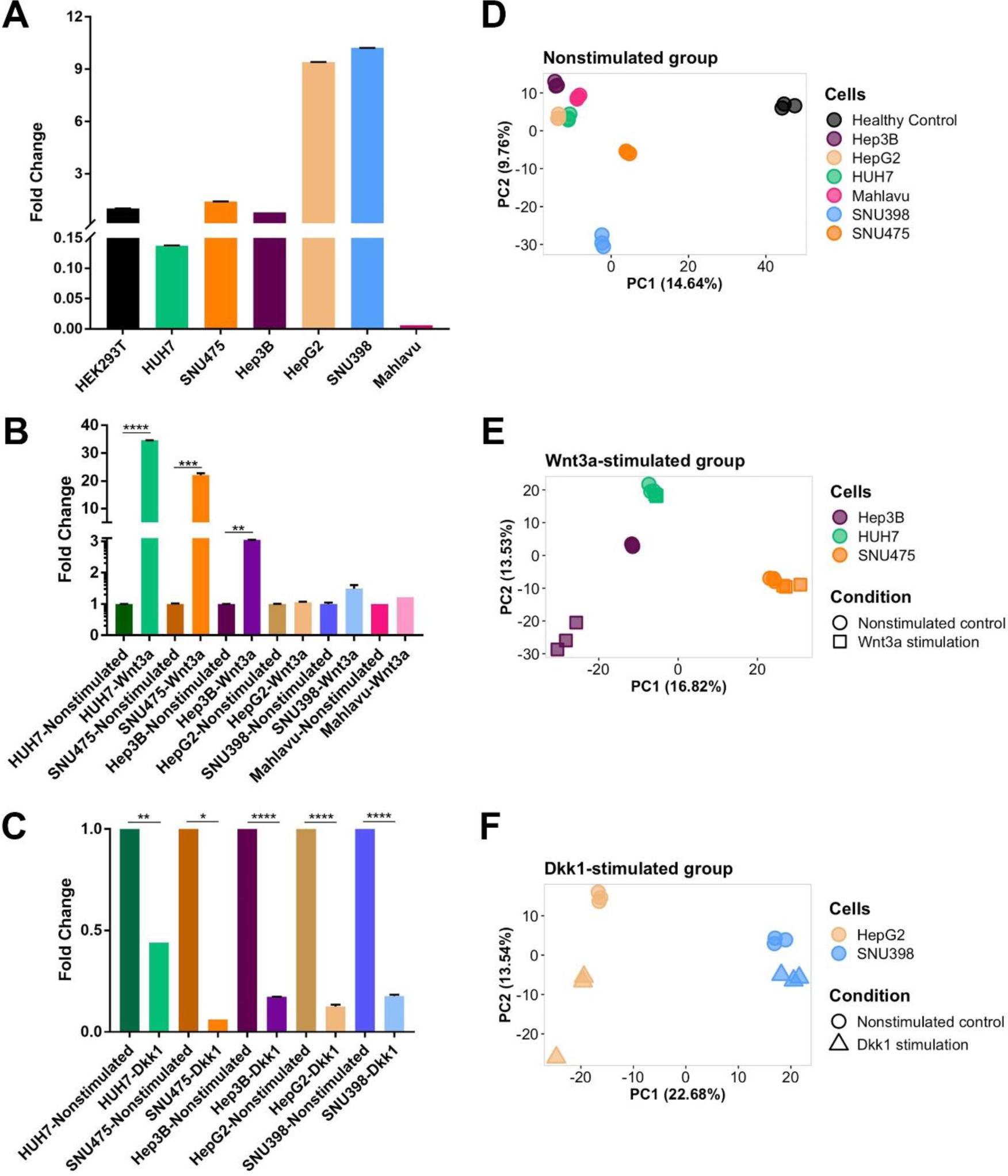
Different HCC cell lines vary in their activity of Wnt/β-catenin signaling and plasma membrane lipid composition. (A-C) Comparion of canonical Wnt signaling activity in various HCC cell lines with known mutations in *β-catenin*, *APC* or *Axin-1* genes. Average and SD of the mean (error bars) values of pBAR luciferase reporter activity represent Wnt/ß-catenin signaling activity (normalized to renilla luciferase activity) in (A) HEK293T, HUH7, SNU475, Hep3B, HepG2, SNU398 and Mahlavu cells, (B) nonstimulated and Wnt3a-stimulated HUH7, SNU475, Hep3B, HepG2, SNU398 and Mahlavu cells and (C) nonstimulated and Dkk1-stimulated HUH7, SNU475, Hep3B, HepG2 and SNU398 cells. Statistical significance was evaluated using unpaired t-test. **** p < 0.0001, *** p < 0.001, ** p < 0.01, and * p < 0.05. Error bars represent SD. Three independent experiments were performed. (D) Principal component analysis (PCA) of nonstimulated healthy control and HCC cells. Cell lines are represented by different colors. (E) PCA of nonstimulated and Wnt3a-stimulated Hep3B, HUH7 and SNU475 cells. (F) PCA of nonstimulated and Dkk1-stimulated HepG2 and Snu398 cells. Cell lines in (E-F) are represented by different colors and stimulation statuses by different shapes.

### Plasma membranes of different HCC cells diverge with respect to distribution of main lipid categories and lipid species

To understand how plasma membrane lipidome profiles alter between different HCC cells, we plotted the log2 fold change of the lipids that were detected in the steady state by mass spectrometry in six different HCC cells as compared to the healthy control liver (THLE2) cells (Figure 2). A great number of glycerophospholipids were differentially regulated in all nonstimulated HCC cell types as compared to the healthy control cells (Figure 2A). Glycerophospholipid intermediates were mostly upregulated (Up) in HCC cells except for SNU475 and Mahlavu cells. Sterols, sphingolipids and glycerolipids were not uniformly altered in one specific direction between HCC and healthy control cells (Figure 2A). Next, we examined the differential regulation patterns with respect to the alterations in the following lipid classes: CE (cholesterol esters), Cer (ceramide), CL (cardiolipin), DAG (diacylglycerol), HexCer (hexosylceramide), LPA (lyso-phosphatidate), LPC (lyso-phosphatidylcholine), LPE (lyso-phosphatidylethanolamine), LPG (lyso-phosphatidylglycerol), LPI (lyso-phosphatidylinositol), LPS (lyso-phosphatidylserine), PA (phosphatidate), PC (phosphatidylcholine), PE (phosphatidylethanolamine), PG (phosphatidylglycerol), PI (phosphatidylinositol), PS (phosphatidylserine), SM (sphingomyelin), TAG (triacylglycerol). When examined in terms of lipid classes, CE, Cer, HexCer and SM (except for Mahlavu cells) were generally downregulated (Down) and LPA, LPC, LPE, LPI, LPS, PI and PS were generally Up in HCC cell lines compared to the healthy control cells (Figure 2B). Notably, most PE class of lipids were Down in SNU398 but Up in HUH7, and several lipids of this class were dramatically Down in Hep3B and Mahlavu cells. In addition, a vast majority of the lipids in the PG and PI classes were Up in all HCC cell lines (Figure 2B). Overall, in accordance with the PCA, differential regulation of lipids is most remarkable in Hep3B, SNU398 and Mahlavu cells with respect to the lipid numbers and statistical significance (Figure 2B).

**Figure 2:**
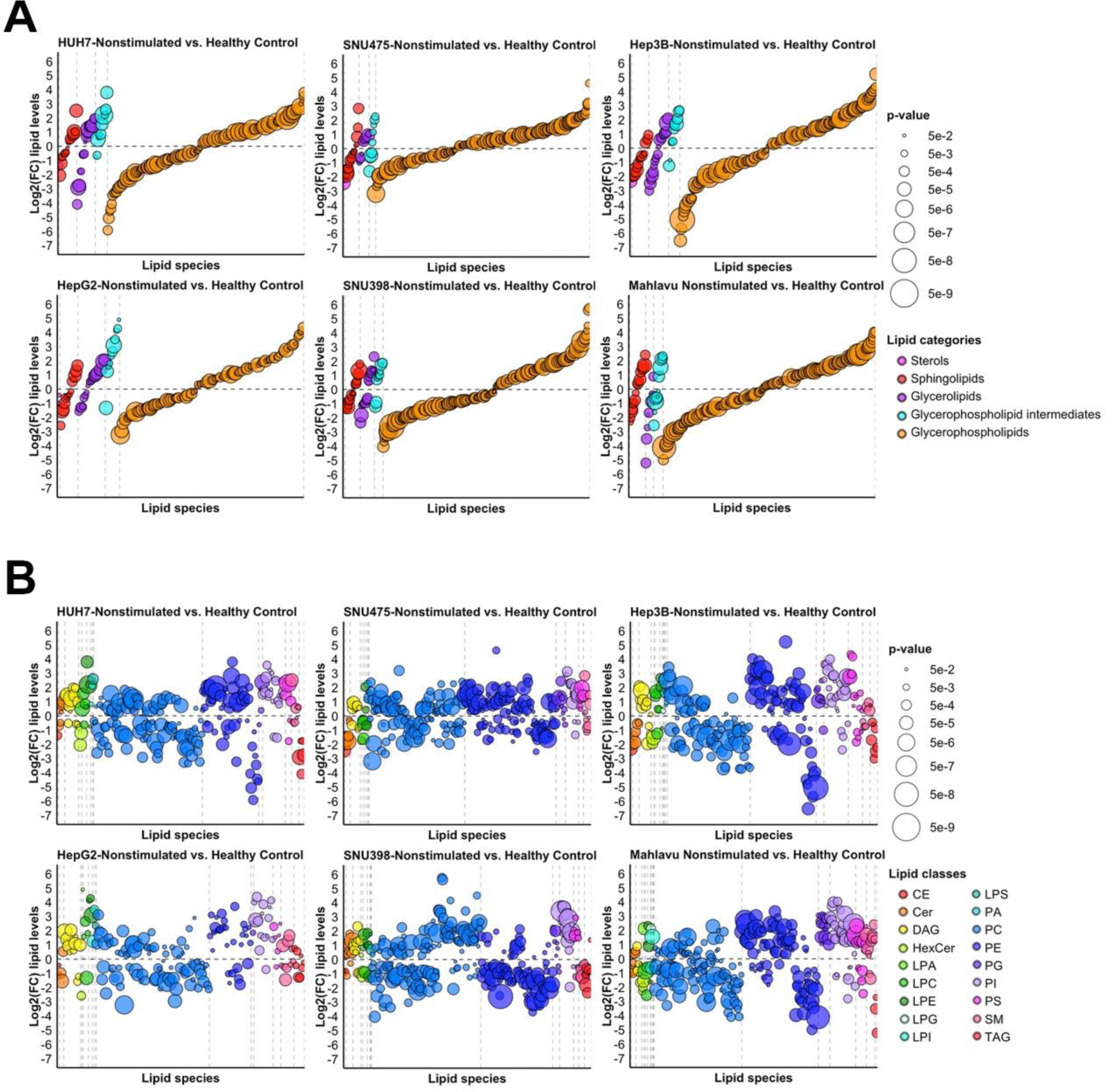
Plasma membranes of different HCC cells diverge with respect to distribution of main lipid categories and lipid species. Differential regulation plots of lipids arranged by (A) lipid categories and (B) lipid classes in nonstimulated HUH7, SNU475, Hep3B, HepG2 and SNU398 cell lines in comparison to healthy control liver cells (THLE2). Lipids are represented by dots arranged regularly on the x-axis with their log2 fold change represented on the y-axis. Dots are colored by (A) lipid category or (B) lipid class and sized proportionally to statistical significance of differential regulation. Only differentially regulated lipid species are shown for each contrast. Vertical dashed lines separate lipid categories/classes. Horizontal dashed line separates up- and down-regulated lipids.

### Membrane lipids of HCC cells alter significantly in response to activation or inhibition of Wnt/β-catenin signaling

Next, we compared the lipidome profiles of the HCC cells that respond to Wnt3a or Dkk1 stimulation to those of the non-stimulated cells by plotting the log2 fold changes of the lipids that are differentially regulated after stimulation (Figure 3, Table S1). Wnt3a induction had a milder effect on all lipid categories in Huh7 and SNU475 cells, while it resulted in strong differential regulation, and more prominently downregulation, of the lipids in Hep3B cells (Figure 3A, Table S1). In HUH7 and SNU475 cells stimulated with Wnt3a, Cer, DAG, PG and PI classes of lipids were mostly Down whereas most lipids in PE class were Up as compared to their nonstimulated counterparts (Figure 3B, Table S1). PC class of lipids appeared to be regulated mildly and oppositely in HUH7-Wnt3a and SNU475-Wnt3a cells. In Hep3B-Wnt3a cells, the number of differentially regulated lipids (DRLs) were much higher and most lipids in the Cer, DAG and LPC classes were more strongly Down (Figure 3B, Table S1). Majority of the lipids in PC, PE, PG, PI, PS and SM classes were likewise Down in Hep3B-Wnt3a cells.

**Figure 3:**
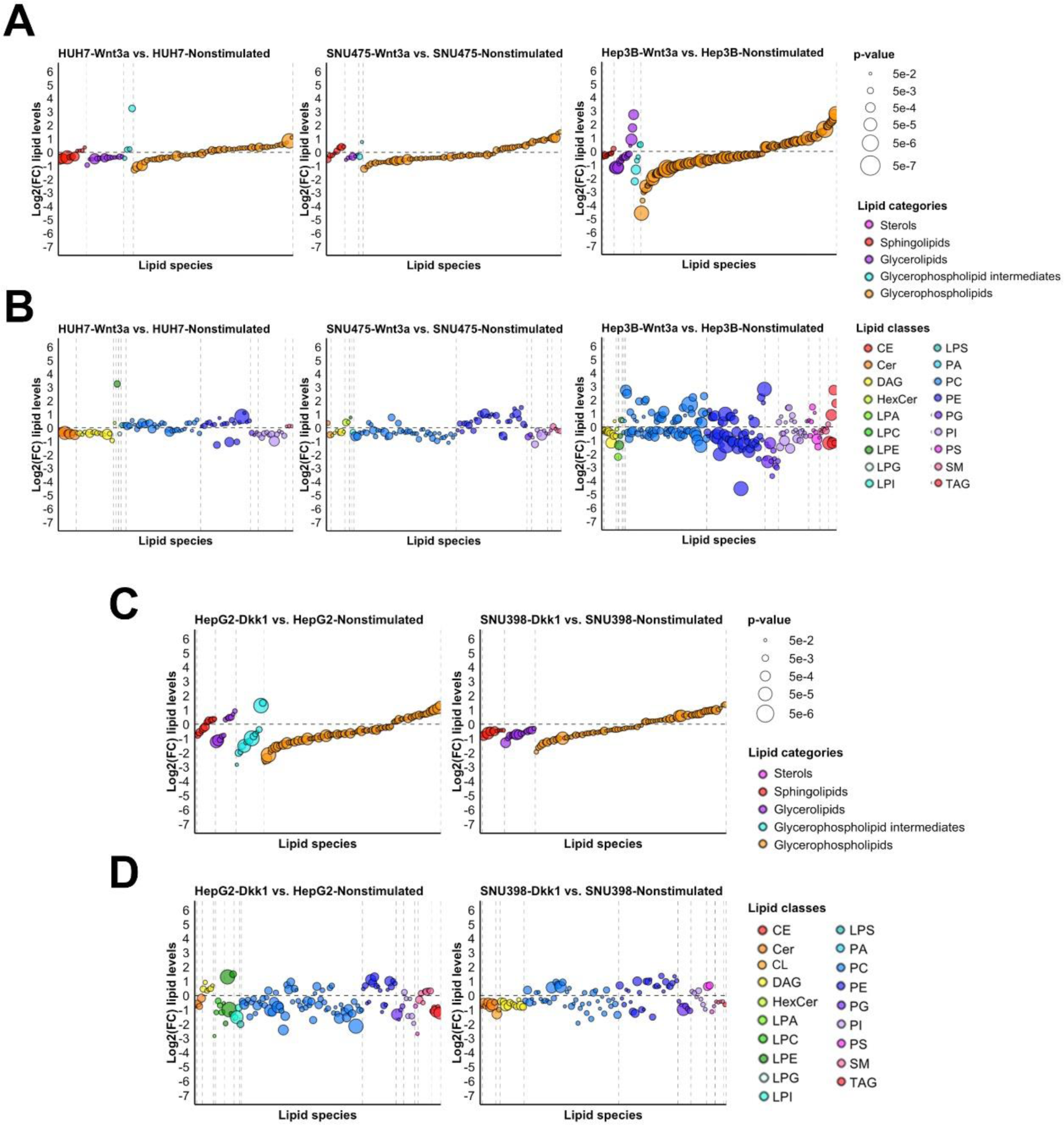
Membrane lipids of HCC cells alter significantly in response to activation or inhibition of Wnt/β-catenin signaling. Differential regulation plots of lipids in response to Wnt signaling pathway manipulation in selected HCC cell lines. (A-B) DRLs arranged by (A) lipid categories and (B) lipid classes in response to Wnt3a stimulation in HUH7, SNU475 and Hep3B cell lines. (C-D) DRLs arranged by (C) lipid categories and (D) lipid classes in response to Dkk1 stimulation in HepG2 and SNU398 lines. Lipids are represented by dots arranged regularly on the x-axis with their log2 fold change represented on the y-axis. Dots are colored by (A, C) lipid category or (B, D) lipid class and sized proportionally to statistical significance of differential regulation. Only differentially regulated lipid species are shown for each contrast. Vertical dashed lines separate lipid categories/classes. Horizontal dashed line separates up- and down-regulated lipids.

Inhibition of Wnt signalling by Dkk1 stimulation resulted in differential regulation of a larger number of lipids in HepG2 cells than in SNU398 cells (Figure 3C, Table S1). In HepG2-Dkk1 cells, the majority of the DRLs, including the glycerophospholipid intermediates, were Down. Interestingly, sphingolipids and glycerolipids were unanimously Down in SNU398-Dkk1 cells while glycerolipids were almost equally distributed as Up and Down. While lipids of the DAG and SM classes were mostly Up in HepG2-Dkk1 cells, they were uniformly Down in SNU398-Dkk1 cells (Figure 3D, Table S1). Cer class of lipids were Down in both HepG2-Dkk1 and SNU398-Dkk1 cells. Although PC class of lipids were overall Down in both cell types after Dkk1 stimulation, some members of this class displayed relatively strong upregulation signals in SNU398-Dkk1 cells. Moreover, lipids in LPA, LPC, LPE and LPI classes showed differential regulation only in HepG2-Dkk1 cells where majority of these lipids were Down (Figure 3D, Table S1). Thus, activation or inhibition of Wnt/β-catenin signaling causes alteration of plasma membrane lipids in HCC cells.

### Global comparison of membrane lipidome profiles reveal differential regulation of lipids in HCC cells and healthy cells

To identify global and specific trends across different HCC cell lines with respect to regulation of lipid classes and categories at the plasma membrane, we pooled all DRLs obtained in comparisons of Wnt3a/Dkk1-stimulated HCC cells to nonstimulated HCC cells as well as of nonstimulated or Wnt3a/Dkk1-stimulated HCC cells to healthy control cells. The generated heatmap summarizes differential regulation profiles of the lipids clustered based on log2 fold changes (Figure 4A). We identified two main clusters of lipid regulation profiles. First, the horizontal clustering of the heatmap showed that the rate of change in lipids observed in HCC cells in response to Wnt pathway manipulation (activation with Wnt3a or inhibition with Dkk1) was much lower than the rate of change in lipids observed in HCC cells compared to healthy cells, independent of manipulation (Figure 4A). Because of the distinct rates of change, lipids were clustered mainly based on the cancer effects rather than the Wnt pathway manipulation effects. In other words, the global influence of “cancer” on the membrane lipidome is greater than that of “Wnt pathway manipulation”. Second, according to the vertical clustering of the heatmap, we identified five main patterns of DRLs (Figure 4A): i) Lipids that were generally Up in HCC cells (with or without Wnt pathway manipulation) as compared to healthy control cells and weakly responsive (mostly as downregulation) to Wnt pathway manipulation, particularly to Wnt3a. ii) Lipids that were generally Down in HCC (with or without Wnt pathway manipulation) as compared to healthy control cells and weakly responsive to Wnt pathway manipulation. iii) Lipids that were specifically Up in SNU398 and SNU475 cells (with or without Wnt pathway manipulation) as compared to healthy control cells. iv) Lipids that were specifically Down in SNU398 and SNU475 cells (with or without Wnt pathway manipulation) as compared to healthy control cells. v) Lipids that were specifically regulated in Hep3B-Wnt3a, HUH7-Wnt3a, HUH7, SNU475-Wnt3a and SNU475 as compared to healthy control cells.

**Figure 4:**
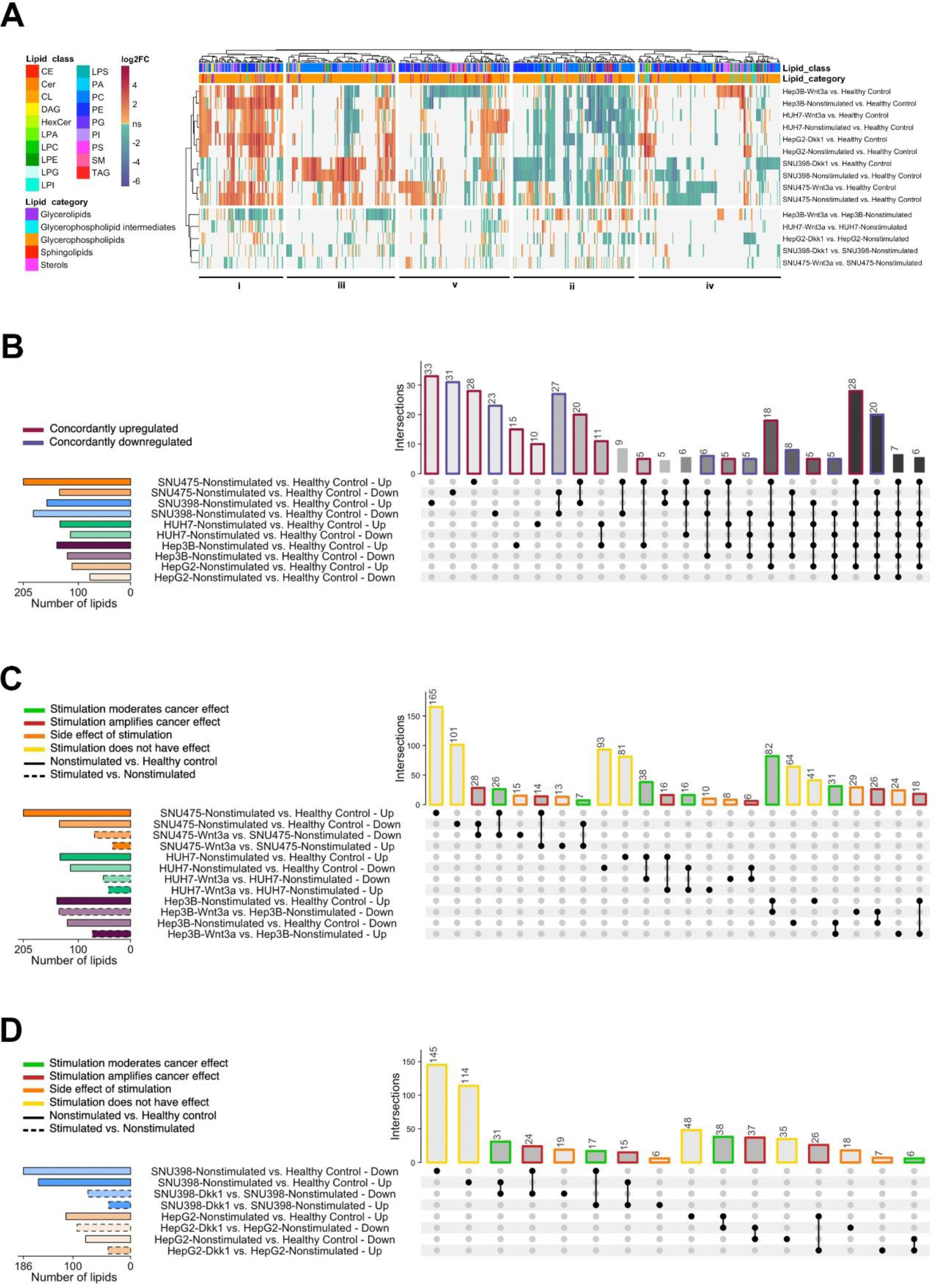
Global comparison of membrane lipidome profiles reveal differential regulation of lipids in HCC cells and healthy cells. (A) Heatmap of lipid species and differential regulation contrasts. Contrasts of the following types are all represented by their differential regulation profiles (log2 fold change): nonstimulated HCC vs. healthy control (cancer effect), stimulated HCC vs. nonstimulated HCC (treatment effect) and stimulated HCC vs healthy control (global effect of cancer and treatment). Contrasts are clustered horizontally into two main blocks: cancer effect and global effect on the upper block and treatment effect on the lower block. Contrasts are clustered vertically into five main blocks indicated as i-v. (B) UpSet plot showing the commonalities of lipid differential regulation between nonstimulated HCC cell lines. The UpSet plot shows the total numbers of DRLs in each cell line on horizontal bars, separated by direction of regulation (Up or Down), as well as the intersections of these sets of lipids on vertical bars. In red are shown the lipid sets that are concordantly Up and in blue are shown the sets of concordantly Down. Only intersections with more than five lipids are displayed. (C-D) UpSet plots showing the effect of (C) Wnt3a stimulation and (D) Dkk1 stimulation on lipids, relatively to the effect of cancer on these lipids. Comparisons are drawn only within each cell line. DRLs are split in four classes for each cell line: Green lipids on which the stimulation moderates the effect of cancer (i.e. downregulates Up lipids or upregulates Down lipids), red lipids on which the stimulation amplifies the effect of cancer (i.e. further upregulates Up lipids or further downregulates Down lipids), yellow lipids that are affected by cancer but on which the stimulation does not have any effect, and orange lipids that were not affected by cancer but are differentially regulated by the stimulation (side effect). Cancer and stimulation related DRL sets are shown by solid and dashed strokes, respectively.

Next, we compared the membrane lipidome profiles of nonstimulated HCC cell lines to those of healthy cell line, considering their direction of regulation. SNU475 and SNU398 cells showed the highest number of DRLs when compared to healthy control (Figure 4B). In these cells, 20 DRLs were concordantly Up and 27 DRLs were concordantly Down. While many DRLs (28 Up and 31 Down in SNU475, 33 Up and 23 Down in SNU398) were specific to the two SNU cell lines, we also detected 48 DRLs (28 Up, 20 Down) that were shared in all five HCC cell lines (Figure 4B).

To assess the effect of Wnt pathway manipulation by Wnt3a and Dkk1 stimulation on DRLs detected in the membrane of HCC cells, we plotted heatmap of lipid species and differential regulation contrasts in the HCC cells treated with Wnt3a or Dkk1 (Figure 4C-D, Table S1). When compared to the healthy control cells, we termed the lipids that were Up or Down in nonstimulated HCC cells as the “cancer effect”. If the regulation direction of such a lipid was reversed after stimulation with Wnt3a or Dkk1 (i.e. from Up to Down or from Down to Up), we termed this influence as “stimulation moderates cancer effect”. If the regulation direction of such a lipid was retained after stimulation with Wnt3a or Dkk1 (i.e. Up & Up or Down & Down), we termed this influence as “stimulation amplifies cancer effect”. There was also a group of lipids termed “stimulation does not have effect” and not affected by Wnt pathway manipulation. The last group of lipids termed “side effect of stimulation” were not affected in cancer but were Up or Down after Wnt pathway manipulation. We found that the membrane lipids of SNU475 and HUH7 cells were largely irresponsive to Wnt3a stimulation (Figure 4C, 266 lipids for SNU475 cells and 174 lipids for HUH7 cells represented in yellow bars). In SNU475 cells, we observed that Wnt3a stimulation amplifies cancer effect (42 lipids represented in red bars) rather than moderating it (33 lipids represented in green bars, Table S2). However, in HUH7 cells, Wnt3a stimulation reduced cancer effect (54 lipids represented in green bars, Table S2) rather than promoting it (22 lipids represented in red bars). In contrast to the SNU475 and HUH7 cells, Wnt3a stimulation mostly mitigated the cancer effect by reversing the differential regulation of lipids in Hep3B cells (Figure 4C, 113 lipids represented in green bars, Table S2). We also noticed that the number of lipids influenced by side effect of stimulation (53 lipids represented in orange bars) or responded to stimulation by amplifying the cancer effect (44 lipids represented in red bars) were higher in Hep3B cells than in SNU475 and HUH7 cells.

The response of SNU398 cells to Dkk1 stimulation was very similar to that of SNU475 cells to Wnt3a stimulation in that their membrane lipids were mostly irresponsive to stimulation (Figure 4D, 259 lipids represented in yellow bars). Apart from that, the moderating, amplifying and side effects of Dkk1 stimulation on cancer effect were comparable in SNU398 cells (48, 39 and 25 lipids represented in green, red and orange bars, respectively). Interestingly, HepG2 cells were more responsive to Dkk1 stimulation than SNU398 cells were, with less irresponsive lipids and more reversed/enhanced lipids (83, 44 and 63 lipids represented in yellow, green and red bars, respectively). Taken together, these results indicate that the membrane lipids are differentially regulated between HCC cells and healthy cells and that certain DRLs further respond to Wnt pathway manipulation by either alleviating or enhancing the cancer effect.

### DAG and ceramide regulate Wnt/β-catenin signaling activity in SNU475 and HepG2 cells

Although the plasma membrane lipidome profiles of the HCC cells are in general significantly different from each other, certain lipids appear to be similarly regulated in response to manipulation of canonical Wnt signaling manipulation, most likely due to their conserved role in regulation of signaling activity at the plasma membrane. Specifically, we found that the levels of diacylglycerol (DAG) and ceramide at the plasma membrane decreased after activation of Wnt/β-catenin signaling with Wnt3a stimulation (Table S1). Both DAG and ceramide have been associated with Wnt signaling in developmental, homeostatic and tumorigenic processes (19,43–45). Binding of the canonical Wnt ligand to the Fz receptor at the plasma membrane causes endocytic internalization of the Wnt-receptor complex and cointernalization of the membrane lipids that are found in the vicinity of the Wnt-receptor complex (17). Thus, the decline in the membrane levels of DAG and ceramide following Wnt3a stimulation of HCC cells suggests that these lipids are involved in formation of Wnt-receptor complex and downstream signaling activity.

To test this hypothesis, we selected two types of HCC cells for the mechanistic studies: i) SNU475 cells that have relatively lower level of endogenous Wnt/β-catenin activity and can be stimulated with Wnt3a, ii) HepG2 cells that have high level of endogenous Wnt/β-catenin activity and cannot be stimulated with Wnt3a (Figure 1A-B). On the one side, we supplied the cells with the DAG or ceramide. On the other side, to deplete DAG and ceramide, we treated the cells with the enzyme DAG kinase (DGK) and the inhibitor of ceramide synthesis myriocin, respectively. Immunofluorescence staining of SNU475 cells revealed a dramatic increase in both total and active phospho-β-catenin (phosphorylated at Ser675 leading to nuclear localization and transcriptional activation) in groups first treated with DAG or ceramide and next stimulated with Wnt3a as compared to the control and Wnt3a-treated groups (Figure 5A). In contrast, depletion of the lipids with DGK or myriocin followed by Wnt3a stimulation led to a decrease in β-catenin levels (Figure 5A). DAG and ceramide significantly increased the activation of the pBAR reporter of Tcf/Lef-mediated transcription in SNU475 cells that were induced with Wnt3a (Figure 5B). Both DGK and myriocin treatment strongly suppressed Wnt3a-induced signaling activity in SNU475 cells (Figure 5C).

**Figure 5:**
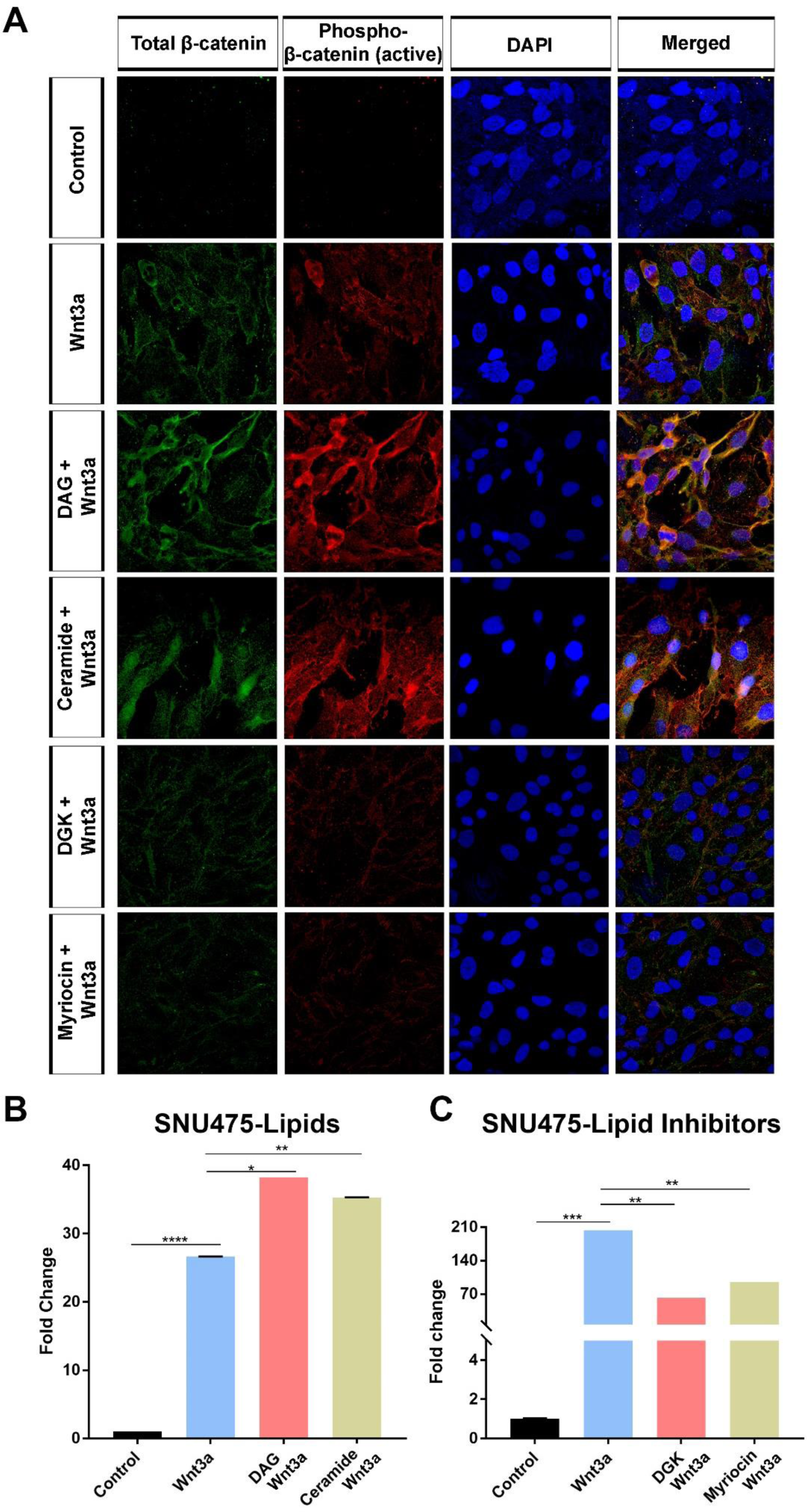
DAG and ceramide regulate Wnt/β-catenin signaling activity in SNU475 cells. (A) Anti total β-catenin (green) and anti-phospho-β-catenin (red) staining of SNU475 cells. Cells are counterstained for DAPI. When compared to Wnt3a-treated cells, treatment of cells with DAG or ceramide increases expression of total and active β-catenin while depletion of DAG with DGK or ceramide with myriocin decreases both types of β-catenin. (B-C) Wnt/ß-catenin signaling activity (normalized to renilla luciferase activity) in Wnt3a-stimulated SNU475 cells treated with (B) DAG or ceramide and (C) DGK or myriocin. Average and SD of the mean (error bars) values of pBAR luciferase reporter activity are shown. Statistical significance was evaluated using unpaired t-test. **** p < 0.0001, *** p < 0.001, ** p < 0.01, and * p < 0.05. Error bars represent SD. Three independent experiments were performed.

In HepG2 cells, depletion of DAG or ceramide reduced endogenous total and active β-catenin (Figure 6A). On the other hand, DAG or ceramide restored the levels of both total and active β-catenin, which were remarkably reduced in response to stimulation with the Wnt inhibitor Dkk1, back to the control level (Figure 6A). DGK and myriocin reduced endogenous canonical Wnt signaling activity in HepG2 cells, at least by 50% (Figure 6B), while DAG rescued the signaling activity that was inhibited by Dkk1 and ceramide treatment resulted in a much stronger activation of Wnt signaling (Figure 6C). Thus, DAG and ceramide are capable of regulating Wnt/β-catenin signaling in HCC cells independently of whether their endogenous signaling activity is low or high.

**Figure 6:**
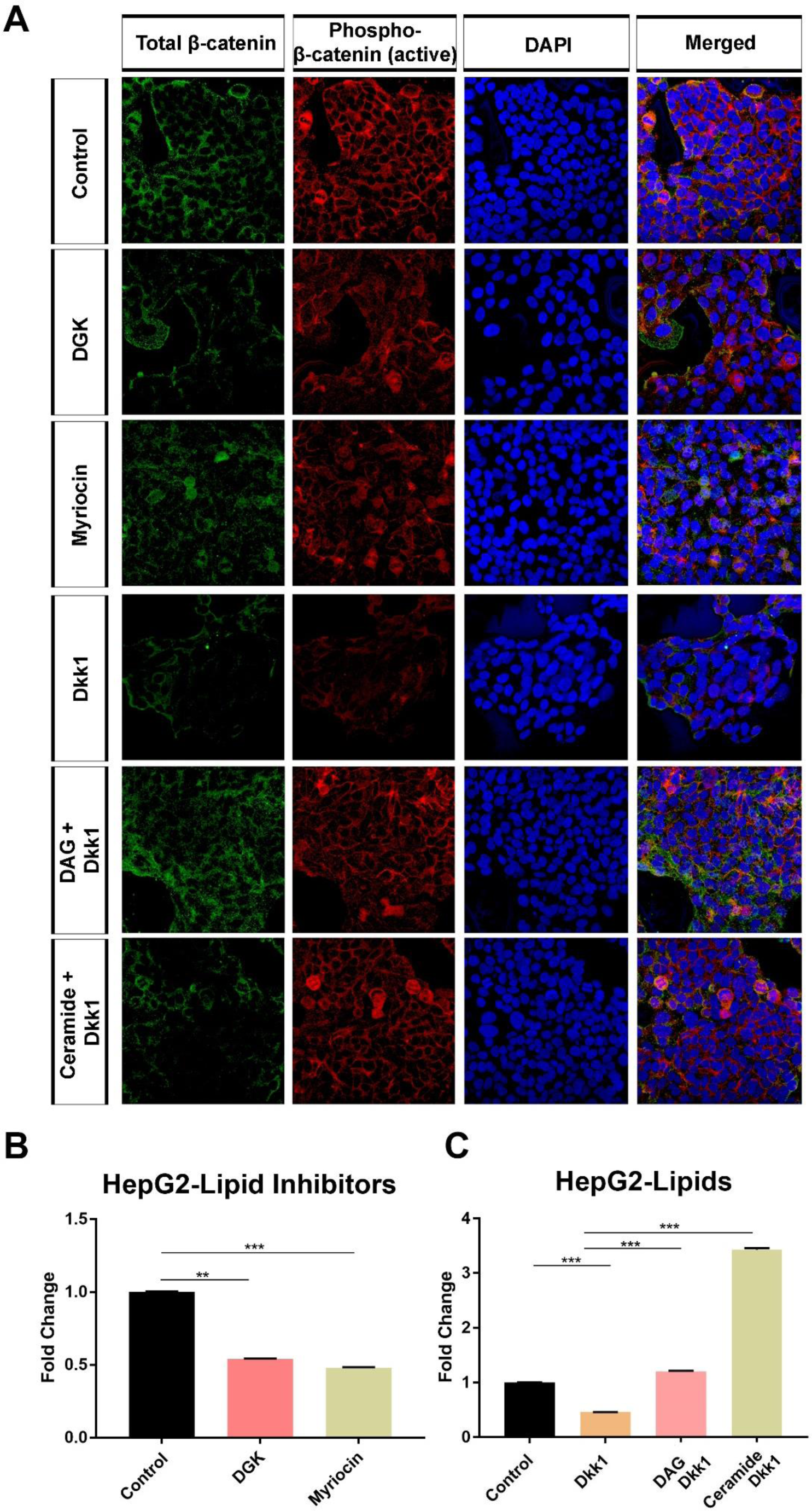
DAG and ceramide regulate Wnt/β-catenin signaling activity in HepG2 cells. (A) Anti total β-catenin (green) and anti-phospho-β-catenin (red) staining of HepG2 cells. Cells are counterstained for DAPI. Depletion of DAG with DGK or ceramide with myriocin decreases expression of total and active β-catenin. Treatment with DAG or ceramide efficiently restores expression of total and active β-catenin, which are potently suppressed by Dkk1. (B-C) Wnt/ß-catenin signaling activity (normalized to renilla luciferase activity) in (B) HepG2 cells treated with DGK or myriocin and (C) Dkk1-stimulated HepG2 cells treated with DAG or ceramide. Average and SD of the mean (error bars) values of pBAR luciferase reporter activity are shown. Statistical significance was evaluated using unpaired t-test. *** p < 0.001 and ** p < 0.01. Error bars represent SD. Three independent experiments were performed.

### Depletion of DAG and ceramide impairs the ability of HCC cells to proliferate and grow into tumors

To examine the effect of DAG and ceramide on HCC cell proliferation, we performed colony formation assay on SNU475 and HepG2 cells. While depletion of DAG with DGK and ceramide with myriocin reduced the colony forming ability of SNU475 cells, a more prominent decrease was detectable when the cells were treated with both lipid synthesis inhibitors (Figure 7A). We observed a similar reduction in the number of colonies formed by HepG2 cells treated with DGK, but not by those treated with myriocin (Figure 7B). Yet, co-treatment of HepG2 cells with DGK and myriocin showed a non-significant reduction in the number of colonies.

**Figure 7:**
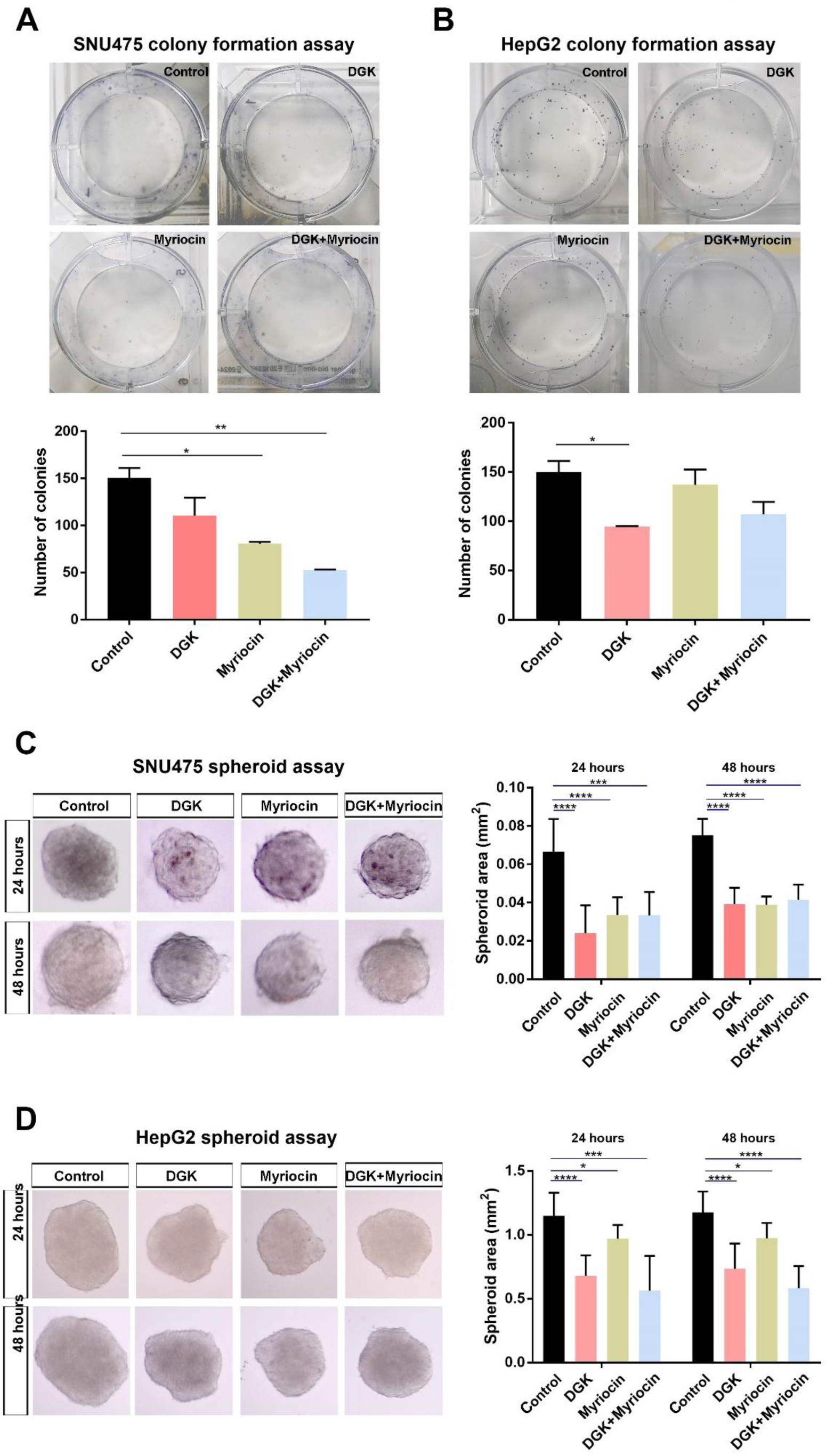
Depletion of DAG and ceramide impairs the ability of HCC cells to proliferate and grow into tumors. (A-B) Colony formation assay and quantification in (A) SNU475 cells and (B) HepG2 cells treated with DGK, myriocin or both. In SNU475 cells, number of colonies significantly decreases after myriocin or combination of inhibitors compared to the control. In HepG2 cells, number of colonies significantly decreases after DGK treatment. (C-D) 3D spheroid formation assay and quantification of spheroid area in (C) SNU475 cells and (D) HepG2 cells treated with DGK, myriocin or both. In SNU475 cells, individual treatment of inhibitors reduce spheroid areas at both 24 and 48 hours while combined treatment does not have an additive effect. In HepG2 cells, individual and combined treatment of inhibitors reduce spheroid areas at both 24 and 48 hours. Statistical significance was evaluated using unpaired t-test. **** p < 0.0001, *** p < 0.001, ** p < 0.01, and * p < 0.05. Error bars represent SD. Three independent experiments were conducted.

Next, to test whether DAG and ceramide are involved in tumor growth, we exploited the spheroid formation assay, which is a scaffold-free 3D model enabling quantitative analysis of tumor growth rate. Individual or combined treatment of SNU475 cells with the lipid synthesis inhibitors DGK and myriocin resulted in significant reduction of the spheroid area at both 24 and 48 hours after addition of the inhibitors (Figure 7C). DGK and myriocin likewise efficiently suppressed the growth of HepG2 spheroids (Figure 7D). These results collectively suggest that reduction of DAG and ceramide efficiently suppresses the proliferation and growth of HCC cells.

### Depletion of DAG and ceramide reduces the migration capacity of HCC cells in vivo

Aberrant regulation of cell migration drives metastasis. To examine the effect of DAG and ceramide on HCC with respect to cell migration and metastasis, we exploited the zebrafish model. We generated larval xenografts by injecting DiI-labeled SNU475 or HepG2 cells into the yolk sac at 2 dpf, treated the xenografts with the lipid synthesis inhibitors and monitored the cancer cell behavior over time. At 4 days post-injection (dpi), activation of canonical Wnt signaling with Wnt3a resulted in increased migration of SNU475 cells as compared to the control (Figure 8A). Individual or combined depletion of DAG with DGK and ceramide with myriocin in Wnt3a-treated SNU475 cells remarkably reduced their migration ability (Figure 8A). DGK and myriocin likewise reduced the migration capacity of HepG2 cells that almost completely invaded the caudal tissue and formed micrometastasis colonies (Figure 8B). The inhibitory effect of DAG or ceramide depletion on HCC cell migration was also reflected in the number of metastatic larvae transplanted with SNU475 cells (Figure 8C) or HepG2 cells (Figure 8D). Thus, depletion of DAG and ceramide can suppress the migratory capacity of HCC cells.

**Figure 8:**
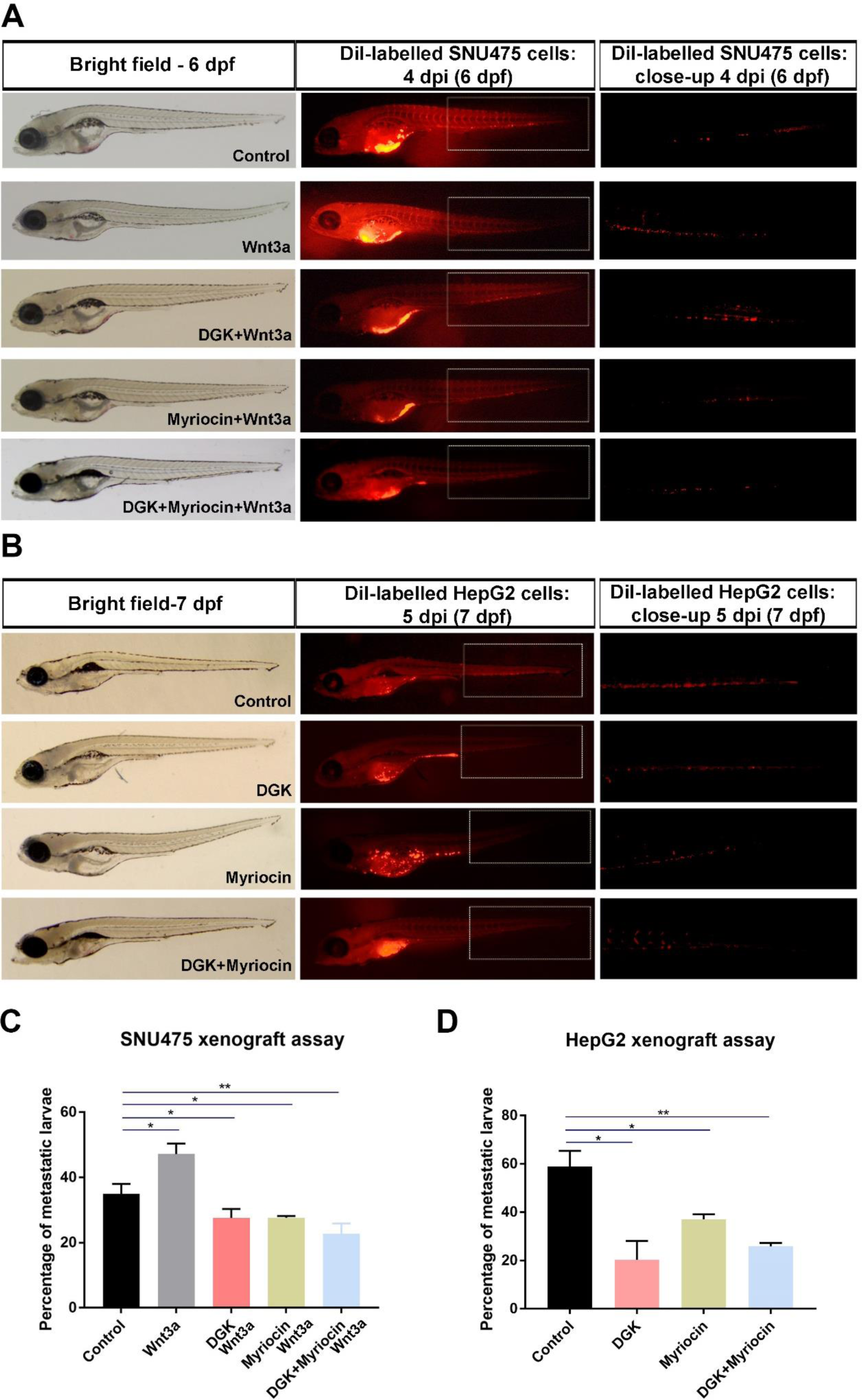
Depletion of DAG and ceramide reduces the migration capacity of HCC cells in vivo. (A) Representative bright-field and fluorescence microscope images of 6 dpf zebrafish embryos injected with SNU475 cells pre-treated with Wnt3a, DGK+Wnt3a, myriocin+Wnt3a or DGK+myriocin+Wnt3a into the yolk sac of 2 dpf zebrafish larvae. While Wnt3a increase the migration capacity of SNU475 cells, individual and combined treatment of inhibitors strongly reduce this capacity even below the control levels. (B) Representative bright-field and fluorescence microscope images of 7 dpf zebrafish embryos injected with HepG2 cells pre-treated with DGK, myriocin or DGK+myriocin into the yolk sac of 2 dpf zebrafish larvae. Individual and combined treatment of inhibitors dramatically reduce the migration capacity of HepG2 cells. (C-D) Quantification of metastatic larvae injected with (C) SNU475 and (D) HepG2 cells. **** p < 0.0001, *** p < 0.001, ** p < 0.01, and * p < 0.05. Error bars represent SD. Three independent experiments were conducted.

## Discussion

Large-scale study of the lipidome, i.e. the complete lipid profile of biological systems, at the quantitative level by mass spectrometry-based lipidomics provides a reasonable approximation of the plasma membrane environment by uncovering membrane lipid composition (34,46,47). Thus, lipidome profiling is a valuable tool to identify the roles of membrane lipids in cellular processes as well as to reveal how lipid composition is altered in cancer. Nevertheless, there is limited information on the lipidome profiles of cancer cells and no data on the comparative analysis of their plasma membrane lipidome profiles. To our knowledge, this is the first study to compare the lipidome profiles of the plasma membranes in different HCC cells. Moreover, we provide the first evidence that manipulation of Wnt/β-catenin signaling dramatically changes the membrane lipid composition in HCC cells. Accordingly, we have reached the following conclusions: i) Six different HCC cells, i.e. HUH7, SNU475, Hep3B, HepG2, SNU398 and Mahlavu cells, vary in their level of endogenous Wnt/β-catenin signaling activity and significantly differ with respect to their plasma membrane lipid composition. ii) HCC cells alter their plasma membrane lipid composition in response to Wnt pathway activation and inhibition. iii) The membrane lipidome profiles of HCC cells differ dramatically from those of healthy cells. There is a considerable amount of lipids that are both differentially regulated in HCC cells and responsive to canonical Wnt pathway manipulation in these cells. iv) DAG and ceramide, which are downregulated after Wnt3a stimulation in HCC cells’ membrane, enhance Wnt/β-catenin signaling activity in SNU475 and HepG2 cells. v) Depletion of DAG and ceramide suppresses Wnt/β-catenin signaling and interferes with the proliferation, tumor formation and *in vivo* migration capacity of SNU475 and HepG2 cells.

By exhibiting aberrant activation of Wnt/β-catenin signaling, HCC would thus constitute a great platform for comparative lipidomics analysis of the plasma membrane (48–50). Comprehensive analysis of the membrane lipidome profiles of HCC cells and healthy control liver cells revealed that while the amount of sterols and sphingolipids were generally lower in the HCC cells than in the healthy cells, glycerophospholipids and their intermediates were more abundant in the HCC cells. Specifically, CE, Cer, HexCer and SM (except Mahlavu) decreased whereas LPA, LPC, LPE, LPI, LPS, PI and PS increased significantly in HCC cell lines compared to the healthy cells. However, we were not able to detect a particular lipidome profile that distinguishes the HCC cells with high endogenous canonical Wnt signaling activity, i.e. HepG2 and SNU398 cells, albeit the overall number of glycerophospholipids were lower in HepG2 cells. Therefore, we decided to examine the response of HCC cells to Wnt signaling manipulation with respect to alteration of the membrane lipids. In general, all lipid categories except for the glycerophospholipids and lipid classes except for PC and PE were downregulated in HUH7, SNU475 and Hep3B cells where canonical Wnt signaling was activated with Wnt3a stimulation. This downregulation influence was most prominent in the levels of Cer, DAG and SM that have previously been associated with the specialized ordered membrane domains, the so-called lipid rafts (51, 52). Ordered membrane domains are dynamically assembled from various saturated lipids, sterols, sphingolipids and lipid-anchored proteins (53). Canonical Wnt ligands, including Wnt3a, bind to their receptors within the ordered domains (17). Canonical Wnt-receptor complex induces recruitment of the cytoplasmic effector Dishevelled to the membrane and formation of the Wnt signalosome (54). A key step in Wnt pathway regulation is internalization of the Wnt signalosome, a critical balance that ensures both pathway activation and endocytic degradation of excessive ligands and receptors (19). Thus, Wnt signalosome internalization inextricably brings about co-internalization of the particular ordered domain lipids that are packed around the Wnt-receptor complex, resulting in their downregulation at the membrane. This outcome is in line with our previous findings where canonical Wnt stimulation reduced the generalized polarization of the membrane, an indicative of reduced membrane order (17).

Wnt3a has been shown to induce caveolin-dependent (clathrin-independent) internalization of the Wnt-receptor complex to activate canonical Wnt signaling, while Dkk1 being associated with clathrin-dependent internalization followed by pathway inhibition (55). Nevertheless, evidence from subsequent studies have reported a role for clathrin also on canonical Wnt signaling by enhancing Wnt signalosome formation and mediating endocytosis of the Wnt-receptor complex (19,54,56,57). Since caveolin-dependent and clathrin-dependent endocytosis take place in the ordered domains, i.e. lipid rafts, and disordered domains, i.e. non-lipid rafts, respectively, stimulation with Wnt3a and Dkk1 are expected to alter the membrane lipid composition differently (58). In HCC cells, when compared to Wnt3a stimulation, Dkk1 stimulation resulted in downregulation of far more lipids in virtually all lipid categories and classes. Owing to its association with the ordered membrane domains, Wnt3a was rather influential on the ordered domains, a restricted portion of the membrane, most likely resulting in a balanced regulation of membrane lipids following internalization. In contrast, due to the expanded nature of clathrin-dependent endocytosis throughout the membrane, Dkk1 stimulation appeared to overwhelm a wider part of the membrane and caused downregulation of a larger number of lipids. It is also noteworthy that stimulation with Wnt3a or Dkk1 resulted in downregulation of Cer in all HCC cells and DAG in all but HepG2-Dkk1 cells. These results suggest that Cer and DAG at the plasma membrane act as common players in regulation of canonical Wnt signaling in response to Wnt3a-mediated activation or Dkk1-mediated inhibition of the pathway.

Comprehensive bioinformatic analyses of membrane lipidome profiles of the Wnt3a/Dkk1-stimulated HCC cells, the nonstimulated HCC cells and the healthy control cells revealed a clear distinction in the clustering patterns of DRLs. Strikingly, this difference was most pronounced in clustering of the DRLs detected in comparison of the HCC cells, regardless of whether they were stimulated or not, to the healthy cells. The difference in DRLs between the stimulated and the nonstimulated HCC cells was rather mild, leading to a weaker clustering. Thus, for the membrane lipid composition of an HCC cell, the influence of being cancerous is stronger than that of having abnormal Wnt signaling activity. There was a considerable number of lipids that was differentially regulated in all HCC cells. When examined further, some of these DRLs were oppositely regulated between cancer and pathway manipulation. In other words, in a particular HCC cell line several lipids were upregulated as compared to the healthy control line and downregulated after manipulation of the Wnt pathway, and several lipids were downregulated compared to the control and upregulated after pathway manipulation. We termed this restoration effect of pathway manipulation as “moderation of cancer by Wnt pathway stimulation” and marked the DRLs that mediate this effect in “green”. Overall, the “green” lipids that were oppositely responsive to cancer formation and Wnt pathway manipulation could be promising targets for the treatment of hepatocellular carcinoma.

Based on the membrane lipidomics analysis, we found that Cer and DAG decreased significantly in response to activation of canonical Wnt pathway in all three HCC cells treated with Wnt3a. Our mechanistic experiments in SNU475 cells (low level of endogenous Wnt/β-catenin activity and can be stimulated with Wnt3a) and HepG2 cells (high level of endogenous Wnt/β-catenin activity and cannot be stimulated with Wnt3a) revealed Cer and DAG were able to enhance Wnt signaling. On the other hand, depletion of Cer and DAG potently inhibited Wnt signaling in these cells and efficiently suppressed their proliferation, tumoral growth and migration capacity. Importantly, Cer and DAG were among the “green” lipids, strongly suggesting that they moderate the cancer effect. The neutral sphingomyelinase 2 enzyme has been shown to induce production of Cer, a sphingolipid component of ordered membrane domains, which induces receptor clustering and plasma membrane curvature in the neural crest during development (52, 59). Subsequent endocytosis of Wnt and BMP signaling complexes was sufficient to activate epithelial-to-mesenchymal transition, suggesting a key role for Cer in regulating cell migration. Thus, the decrease in Cer in Wnt3a-stimulated HCC cells is likely due to its endocytosis with the Wnt-receptor complex. In a different study, ceramides have been proposed to increase the level of exosomal miRNAs that activate Wnt/β-catenin signaling and enhance cell invasion (60). This is in line with the inhibition of cell migration upon depletion of Cer in SNU475 and HepG2 cells. Furthermore, ceramide metabolism has been reported to be dysregulated in HCC (61–63). DAG is a key signaling molecule that has been identified as a component of the ordered membrane domains (64–66). Elevated levels of DAG promotes colony-stimulating factor 1-dependent proliferation in DGK knock-out mice and modulates β-catenin/cyclinD1 levels in osteoclast precursors (67). DAG has also been associated with HCC progression (68, 69). DAG, by either DGK-mediated phosphorylation into phosphatidic acid or activating protein kinase C, supports tumor growth and HCC progression (69–71). We observed a parallel response to DAG in SNU475 and HepG2 cell lines. Therefore, Cer and DAG harbors a potential for diagnosis and targeted therapy in patients with HCC, especially when Wnt/β-catenin signaling is aberrantly activated.

Liver lipidome has been examined in tissue and serum samples of patients with HCC or non-alcoholic fatty liver disease (NAFLD), a chronic liver disease and the fastest growing cause of HCC (72–76). Due to liver’s essential roles in lipid metabolism, altered lipid metabolism in hepatic cells contributes to the progression of HCC, by providing the neoplastic cells with energy and supporting the growth of HCC lesions in humans (77). The changes in the lipid metabolism and the levels of different lipids have been associated with the severity of HCC (74). However, these studies are mainly based on the data obtained from whole tissues or serum, presumably leading to a heterogeneity in the role of the lipids in HCC. As our data reveal that various lipids, including Cer and DAG, are differentially regulated between different types of HCC, it is also very likely that their functions vary depending on the cancer subtype. Thus, membrane lipidome analysis is a key aspect for individual and cell type-dependent identification and characterization of the lipids in cancer.

## Conclusion

Lipids are essential components of the plasma membrane that is known to alter significantly in cancer cells as compared to the normal cells. Comparative lipidome profiling of the plasma membranes of cancer cells and healthy cells can greatly contribute to our understanding of the role of specific membrane lipids in the onset and progression of cancer and to identification of specific lipids including Cer and DAG that could serve as effective diagnostic or prognostic biomarkers and therapeutic targets in cancer.

## Acknowledgement

We would like to thank Prof. Randall Moon (University of Washington) for the plasmid pCS2P+dkk1GFP. We would like to thank Izmir Biomedicine and Genome Center Vivarium-Zebrafish Core Facility and the facility manager Emine Gelinci, Optical Imaging Core Facility and the facility manager Dr. Melek Ucuncu and Histopathology Core Facility for providing zebrafish care, microscope facility support and histopathology service support, respectively. We would like to thank Berfin Yesil for her assistance. We also thank Lipotype GmbH, Dresden, Germany for the lipidomics analysis.

## Ethics Statement

The animal study was reviewed and approved by the Animal Experiments Local Ethics Committee of Izmir Biomedicine and Genome Center (IBG-AELEC).

## Author Contributions

GO and YA designed the experiments. YA and MK performed the molecular and cell biology experiments. YD and GH conducted the bioinformatics analyses. YA drafted the manuscript, YD and GH contributed to the results, and materials and methods. GO wrote the manuscript.

